# Antibody Resistance of SARS-CoV-2 Variants B.1.351 and B.1.1.7

**DOI:** 10.1101/2021.01.25.428137

**Authors:** Pengfei Wang, Manoj S. Nair, Lihong Liu, Sho Iketani, Yang Luo, Yicheng Guo, Maple Wang, Jian Yu, Baoshan Zhang, Peter D. Kwong, Barney S. Graham, John R. Mascola, Jennifer Y. Chang, Michael T. Yin, Magdalena Sobieszczyk, Christos A. Kyratsous, Lawrence Shapiro, Zizhang Sheng, Yaoxing Huang, David D. Ho

## Abstract

The COVID-19 pandemic has ravaged the globe, and its causative agent, SARS-CoV-2, continues to rage. Prospects of ending this pandemic rest on the development of effective interventions. Single and combination monoclonal antibody (mAb) therapeutics have received emergency use authorization^1–3^, with more in the pipeline^4–7^. Furthermore, multiple vaccine constructs have shown promise^8^, including two with ~95% protective efficacy against COVID-19^9,10^. However, these interventions were directed toward the initial SARS-CoV-2 that emerged in 2019. The recent emergence of new SARS-CoV-2 variants B.1.1.7 in the UK^11^ and B.1.351 in South Africa^12^ is of concern because of their purported ease of transmission and extensive mutations in the spike protein. We now report that B.1.1.7 is refractory to neutralization by most mAbs to the N-terminal domain (NTD) of spike and relatively resistant to a few mAbs to the receptor-binding domain (RBD). It is not more resistant to convalescent plasma or vaccinee sera. Findings on B.1.351 are more worrisome in that this variant is not only refractory to neutralization by most NTD mAbs but also by multiple individual mAbs to the receptor-binding motif on RBD, largely due to an E484K mutation. Moreover, B.1.351 is markedly more resistant to neutralization by convalescent plasma (9.4 fold) and vaccinee sera (10.3-12.4 fold). B.1.351 and emergent variants^13,14^ with similar spike mutations present new challenges for mAb therapy and threaten the protective efficacy of current vaccines.

---

Considerable SARS-CoV-2 evolution has occurred since its initial emergence, including variants with a D614G mutation^15^ that have become dominant. Viruses with this mutation alone do not appear to be antigenically distinct, however^16^. SARS-CoV-2 B.1.1.7, also known as 501Y.V1 in the GR clade (Fig. 1a), emerged in September 2020 in South East England and rapidly became the dominant variant in the UK, possibly due to its enhanced transmissibility^11^. This strain has now spread to over 50 countries, and there are indications that it may be more virulent^17^. B.1.1.7 contains 8 spike mutations in addition to D614G, including two deletions (69-70del & 144del) in NTD, one mutation (N501Y) in RBD, and one mutation (P681H) near the furin cleavage site (Fig. 1b). SARS-CoV-2 B.1.351, also known as 501Y.V2 in the GH clade (Fig. 1a), emerged in late 2020 in Eastern Cape, South Africa (SA)^12^. This variant has since become dominant locally, raising the specter that it too has enhanced transmissibility. B.1.351 contains 9 spike mutations in addition to D614G, including a cluster of mutations (e.g., 242-244del & R246I) in NTD, three mutations (K417N, E484K, & N501Y) in RBD, and one mutation (A701V) near the furin cleavage site (Fig. 1b). There is a growing concern that these new variants could impair the efficacy of current mAb therapies or vaccines, because many of the mutations reside in the antigenic supersite in NTD^18,19^ or in the ACE2-binding site (also known as the receptor-binding motif—RBM) that is a major target of potent virus-neutralizing antibodies. We therefore addressed this concern by assessing the susceptibility of authentic B.1.1.7 and B.1.351 viruses to neutralization by 30 mAbs, 20 convalescent plasma, and 22 vaccinee sera. In addition, we created VSV-based SARS-CoV-2 pseudoviruses that contain each of the individual mutations as well as one with all 8 mutations of the B.1.1.7 variant (UKΔ8) and another with all 9 mutations of the B.1.351 variant (SAΔ9). A total of 18 mutant pseudoviruses were made as previously described^20,21^, and each was found to have a robust titer (Extended Data Fig. 1) adequate for neutralization studies.

**Fig. 1.**
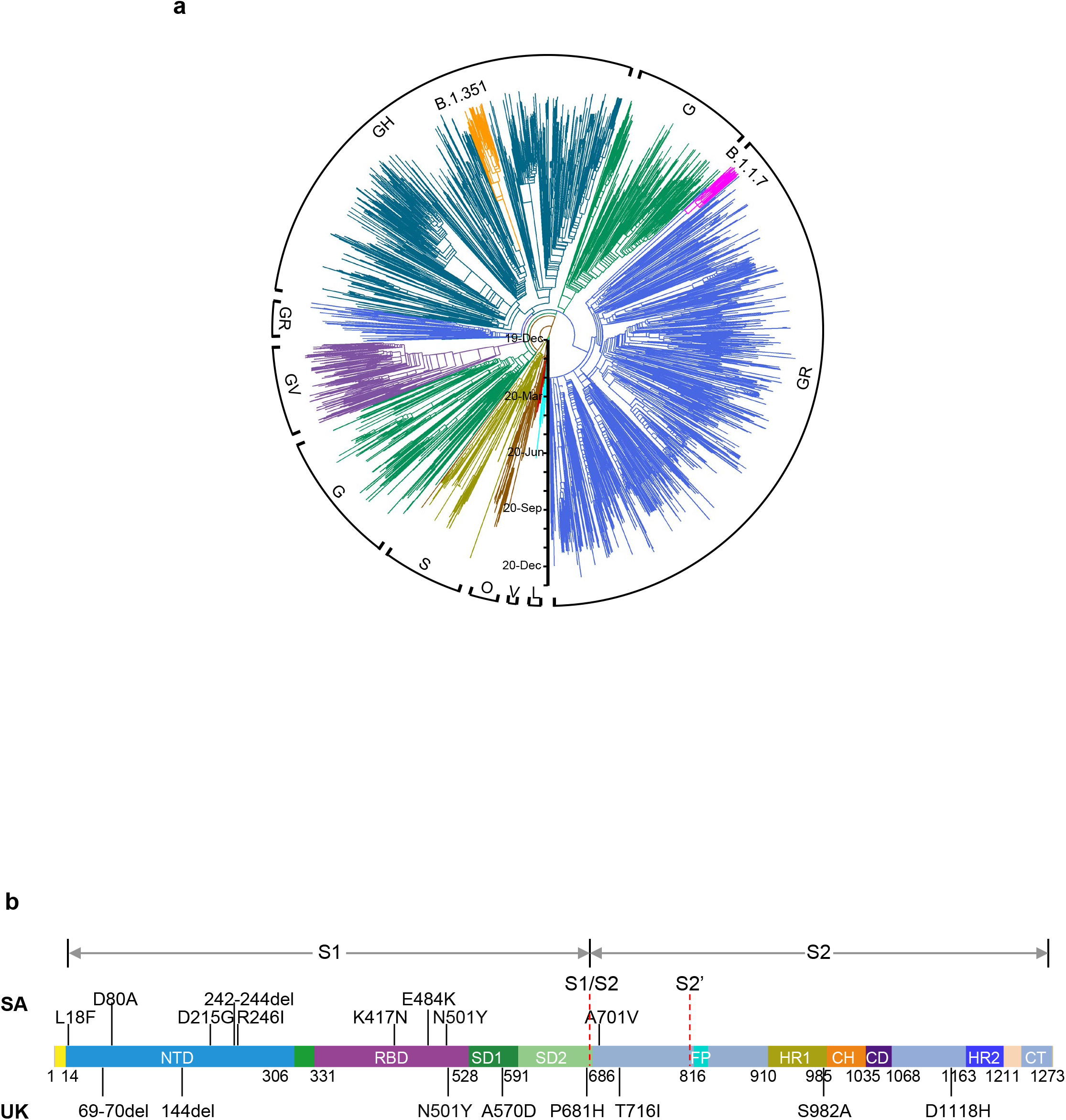
Emerging SARS-CoV-2 variants identified in the United Kingdom and South Africa. **a,** Phylogenetic tree of SARS-CoV-2 variants, with B.1.351 and B.1.1.7 highlighted. **b,** Mutations in the viral spike identified in B.1.351 (SA) and B.1.1.7 (UK) in addition to D614G.

## Monoclonal antibodies

We first assayed the neutralizing activity of 12 RBD mAbs against authentic B.1.1.7 and B.1.351 viruses, as compared to the original SARS-CoV-2 strain (WT), in Vero E6 cells as previously described^20,21^. Three mAbs target the “inner side”, four target RBM, and five target the “outer side”. The footprints of these mAbs on RBD are shown in Fig. 2a, and their neutralization profiles are shown in Fig. 2b. For neutralization of B.1.1.7, only the activities of 910-30^22^ and S309^5^ are significantly impaired. For neutralization of B.1.351, however, the activities of 910-30, 2-15^20^, LY-CoV555 (bamlanivimab)^1,23^, C121^24^, and REGN10933 (casirivimab)^2^ are completely or markedly abolished. The four mAbs that target RBM are among the most potent SARS-CoV-2-neutralizing antibodies in clinical use or development. Note that mAbs directed to lower aspects of the “inner side” (2-36^20^ & COVA1-16^25,26^) or to the “outer side” retain their activities against B.1.351, including 2-7^20^, REGN10987 (imdevimab)^2^, C135^24^, and S309 that are in clinical use or development. The results on the neutralization of B.1.1.7 and B.1.351 by these 12 mAbs are summarized in Fig. 2c as fold increase or decrease in IC50 neutralization titers relative to the WT. To understand the specific spike mutations responsible for the observed changes, we also tested the same panel of mAbs against pseudoviruses UKΔ8 and SAΔ9, as well as those containing only a single mutation found in B.1.1.7 or B.1.351. The results are displayed, among others, in Extended Data Fig. 2 and summarized in Fig. 2c. There is general agreement for results between B.1.1.7 and UKΔ8, as well as between B.1.351 and SAΔ9. Against B.1.1.7, the decreased activity of 910-30 is mediated by N501Y, whereas the slightly impaired activity of S309 is unexplained. Against B.1.351, the complete loss of activity of 2-15, LY-CoV555, and C121 is mediated by E484K; the complete loss for 910-30 is mediated by K417N; and the marked reduction for REGN10933 is mediated by K417N and E484K, as has been reported^27^. A structural explanation on how E484K disrupts the binding of 2-15, LY-CoV555, and REGN10933 is presented in Extended Data Fig. 3a.

**Fig. 2.**
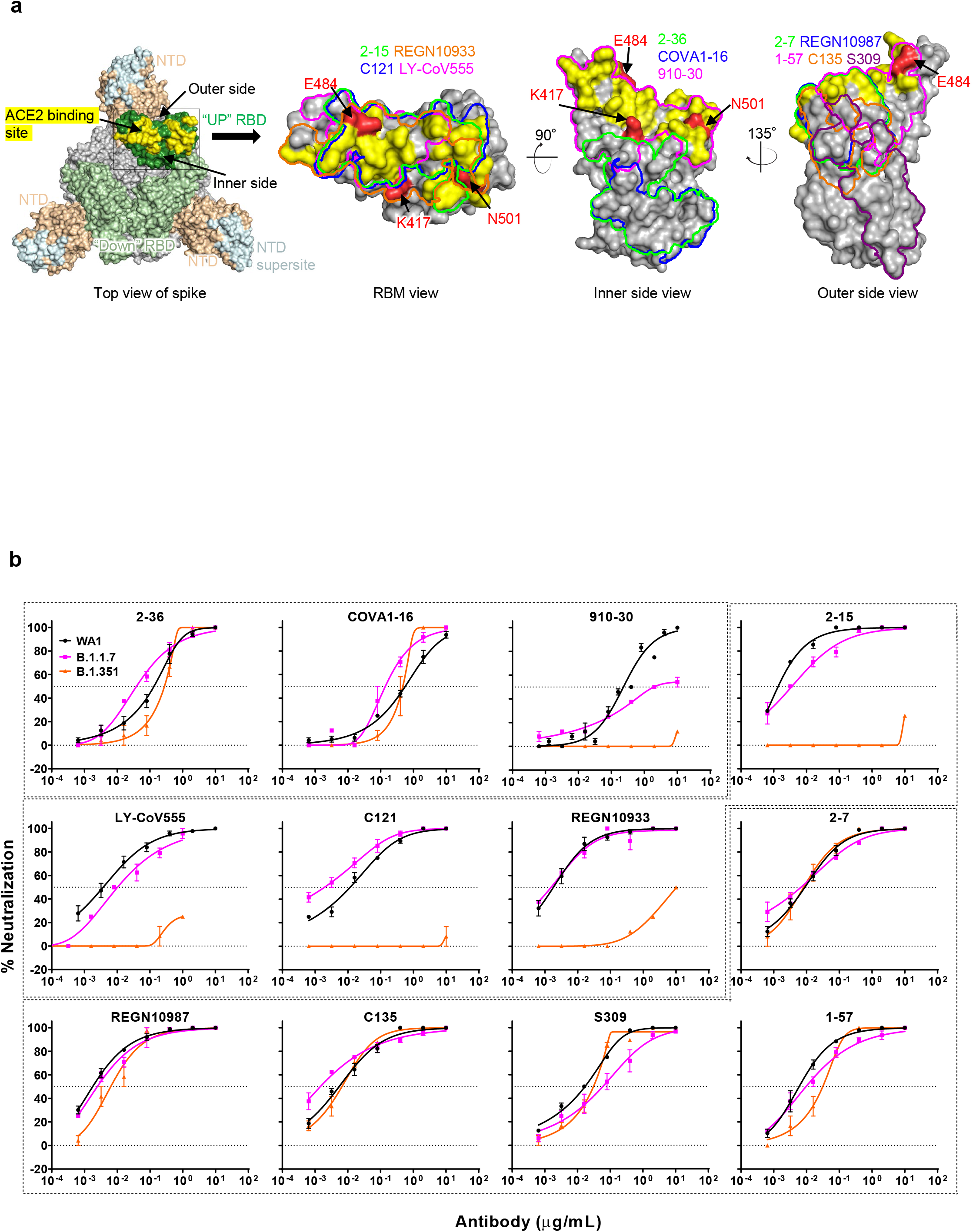
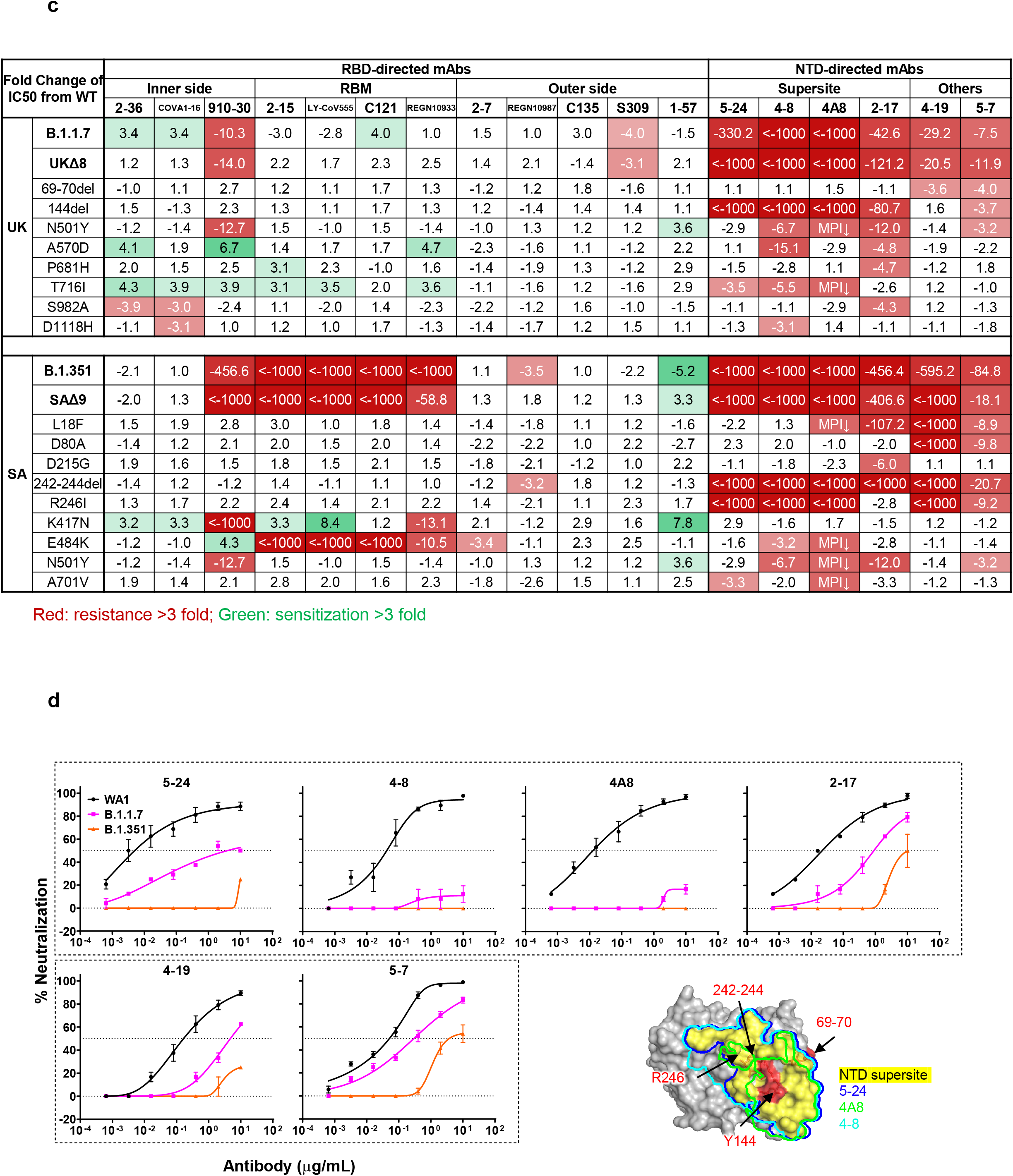
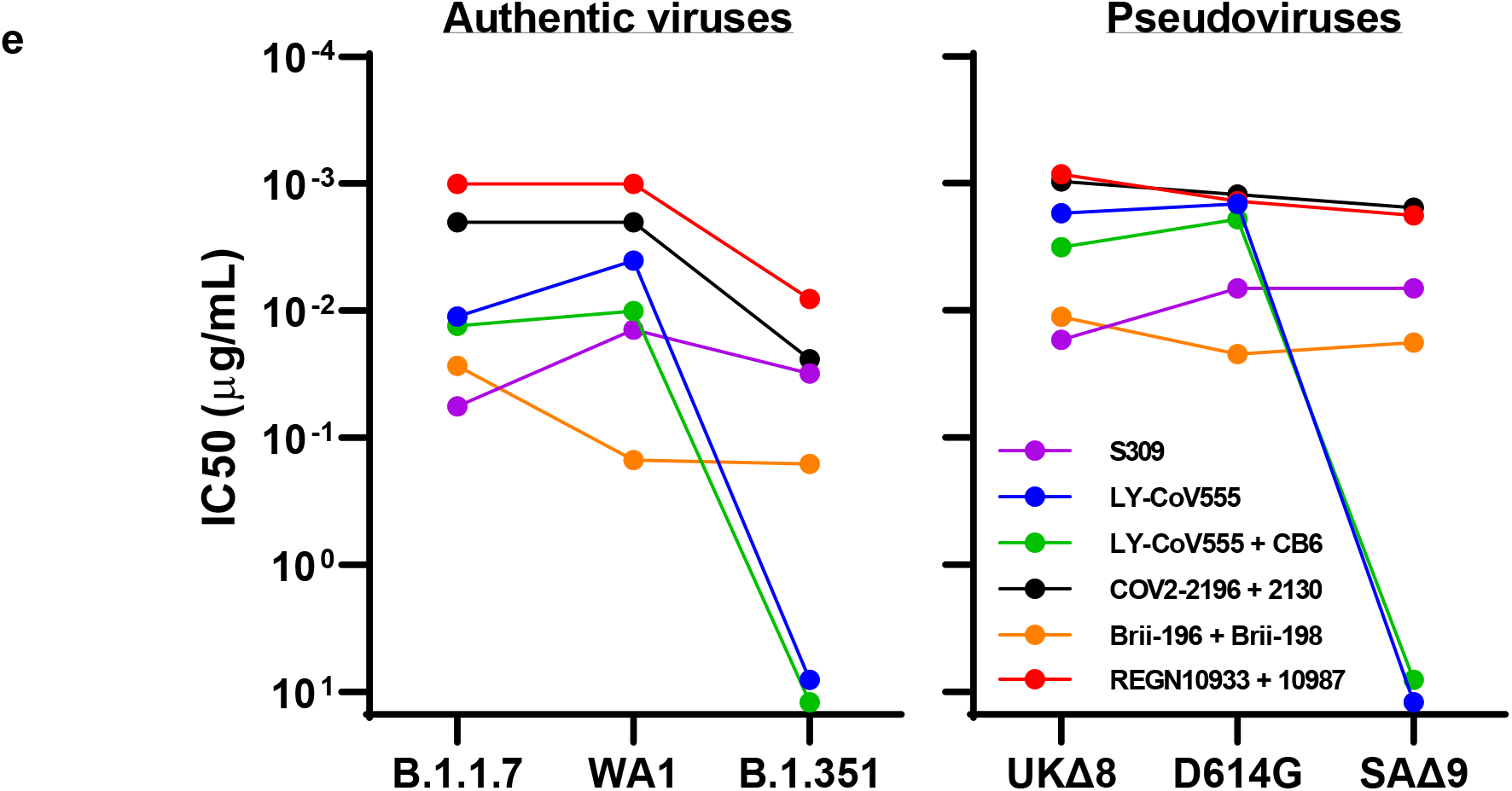
Susceptibility of B.1.1.7 and B.1.351 to neutralization by mAbs. **a,** Footprints of neutralizing mAbs on the RBD. Left panel, top view of SARS-COV-2 spike with one RBD in the “up” conformation (pdb: 6zgg). RBD and NTD are colored green and peach, respectively. The positions of ‘inner’ and ‘outer’ sides are indicated on the “up” RBD with the ACE2-binding site colored yellow. The three panels to the right show the antibody footprints on RBD. **b,** Neutralization of B.1.1.7, B.1.351, and WT viruses by select RBD mAbs. **c,** Fold increase or decrease in IC50 of neutralizing mAbs against B.1.1.7 and B.1.351, as well as UKΔ8, SAΔ9, and single-mutation pseudoviruses, relative to WT, presented as a heatmap with darker colors implying greater change. MPI↓ denotes that maximum percent inhibition is substantially reduced, confounding IC50 calculations. **d,** Neutralization of B.1.1.7, B.1.351, and WT viruses by NTD-directed mAbs, the footprints of which are delineated by the color tracings in the insert. **e,** Changes in neutralization IC50 of authorized or investigational therapeutic mAbs against B.1.1.7, B.1.351, WT (WA1) viruses as well as UKΔ8, SAΔ9, and WT (D614G) pseudoviruses. Data in **b** and **d** are mean ± SEM of technical triplicates, and represent one of two independent experiments.

We also assessed the neutralizing activity of six NTD mAbs against B.1.1.7, B.1.351, and WT viruses. Both B.1.1.7 and B.1.351 are profoundly resistant to neutralization by our antibodies 5-24 and 4-8^20^, as well as by 4A8^28^, all of which target the antigenic supersite in NTD^18^ (Insert in Fig. 2d). The activities of 2-17, 4-19, and 5-7^20^ are variably impaired, particularly against B.1.351. To understand the specific mutations responsible for the observed changes, we then tested these mAbs against pseudoviruses containing only a single mutation found in B.1.1.7 or B.1.351 (Extended Data Fig. 2). The results are summarized in Fig. 2c as fold increase or decrease relative to the WT (D614G). It is evident that the resistance of B.1.1.7 to most NTD mAbs is largely conferred by 144del, whereas the resistance of B.1.351 is largely conferred by 242-244del and/or R246I. Amino-acid residues 144, 242-244, and 246 all fall within the NTD supersite^18,19^ (Insert in Fig. 2d; details in Extended Data Fig. 3b).

We next tested the neutralizing activity of 12 additional RBD mAbs, including ones from our own collection (1-20, 4-20, 2-4, 2-43, 2-30, & 2-38)^20^ as well as CB6 (etesevimab)^3,6^, COV2-2196 & COV2-2130^7^, Brii-196 & Brii-198^4^, and REGN10985. The results against B.1.1.7, B.1.351, and WT are highlighted in Extended Data Fig. 4a, and the detailed findings against the single-mutation pseudoviruses are shown in Extended Data Fig. 2. The fold changes in neutralization IC50 titers relative to the WT are tabulated in Extended Data Fig. 4b. Here, we only comment on results for mAbs in clinical development. The activity of CB6 is rendered inactive against B.1.351 because of K417N. Brii-196 and COV2-2130 are essentially unaffected by the new variants; the activities of Brii-198 and COV2-2196 are diminished 14.6 fold and 6.3 fold, respectively, against B.1.351 but not against B.1.1.7.

Lastly, we examined, in a single experiment, the neutralizing activity of mAb therapies in clinical use or under clinical investigation against B.1.1.7, B.1.351, and WT viruses, as well as against UKΔ8, SAΔ9, and WT pseudoviruses. The results for single mAb LY-CoV555 and S309, as well as for combination regimens REGN10933+REGN10987, LY-CoV555+CB6, Brii-196+Brii-198, and COV2-2196+COV2-2130, are shown in Extended Data Fig. 5 and summarized in Fig. 2e. Note that LY-CoV555, alone or in combination with CB6, is no longer able to neutralize B.1.351. While REGN10933+REGN10987 and COV2-2196+COV2-2130 are seemingly unaffected against variant pseudoviruses, there are noticeable decreases in their activity against B.1.351 authentic virus. Although S309 and the Brii-196+Brii-198 combination are not significantly impaired, their potencies are noticeably lower (Fig. 2e). These findings suggest that antibody treatment of this virus might need to be modified in localities where B.1.351 and related variants^13,14^ are prevalent, and highlight the importance of combination antibody therapy to address the expanding antigenic diversity of SARS-CoV-2.

## Convalescent plasma

We obtained convalescent plasma from 20 patients more than one month after documented SARS-CoV-2 infection in the Spring of 2020. Each plasma sample was then assayed for neutralization against B.1.1.7, B.1.351, and WT viruses. Fig. 3a shows that most (16 of 20) plasma samples lost >2.5-fold neutralizing activity against B.1.351, while maintaining activity against B.1.1.7. Only plasma from P7, P10, P18, and P20 retain neutralizing activities similar to those against the WT. These results are summarized as fold increase or decrease in plasma neutralization IC50 titers in Fig. 3b. Furthermore, the magnitude of the drop in plasma neutralization is better seen in Fig. 3c, showing no loss of activity against B.1.1.7 but substantial loss against B.1.351 (9.4 fold).

**Fig. 3.**
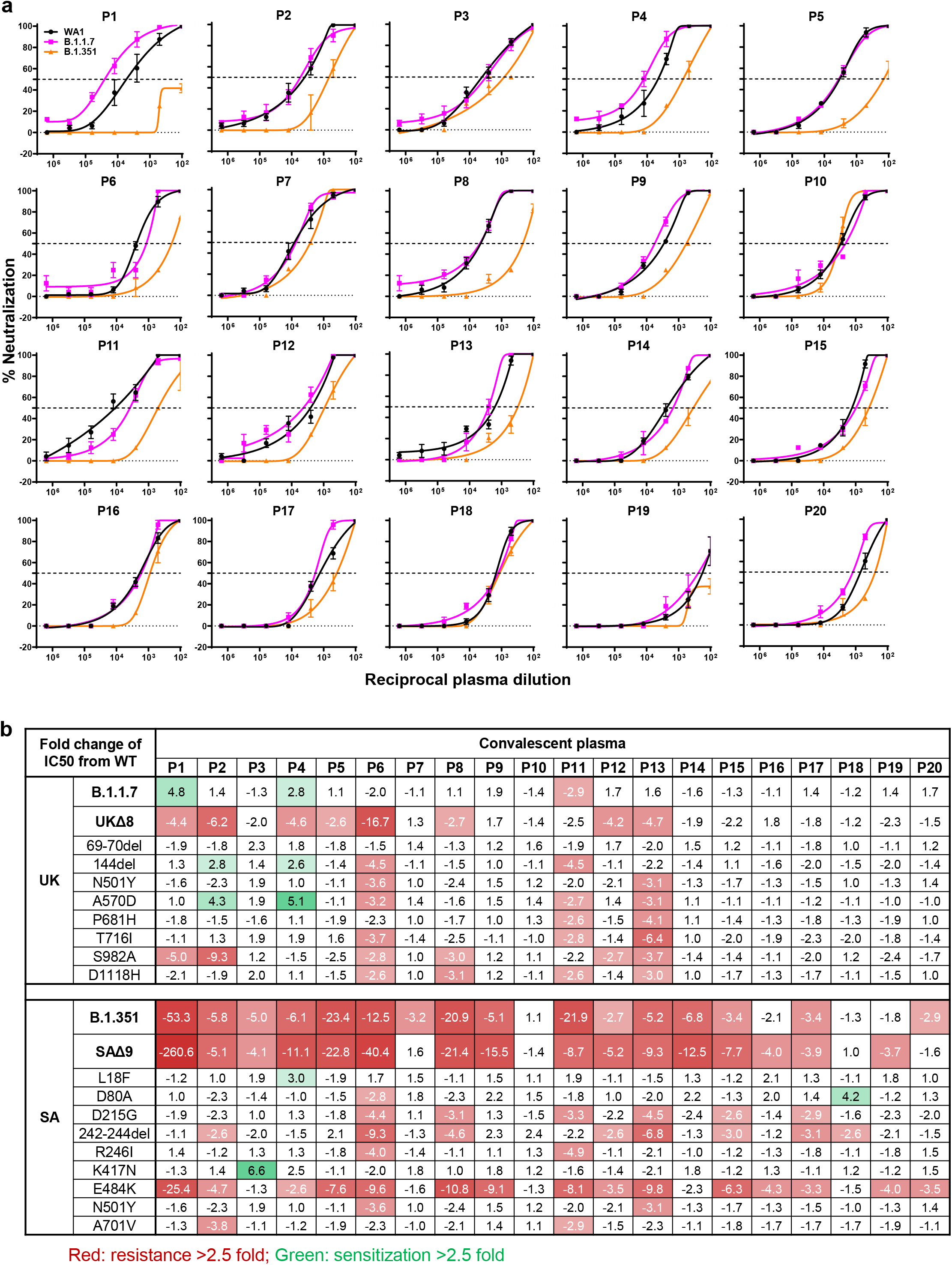
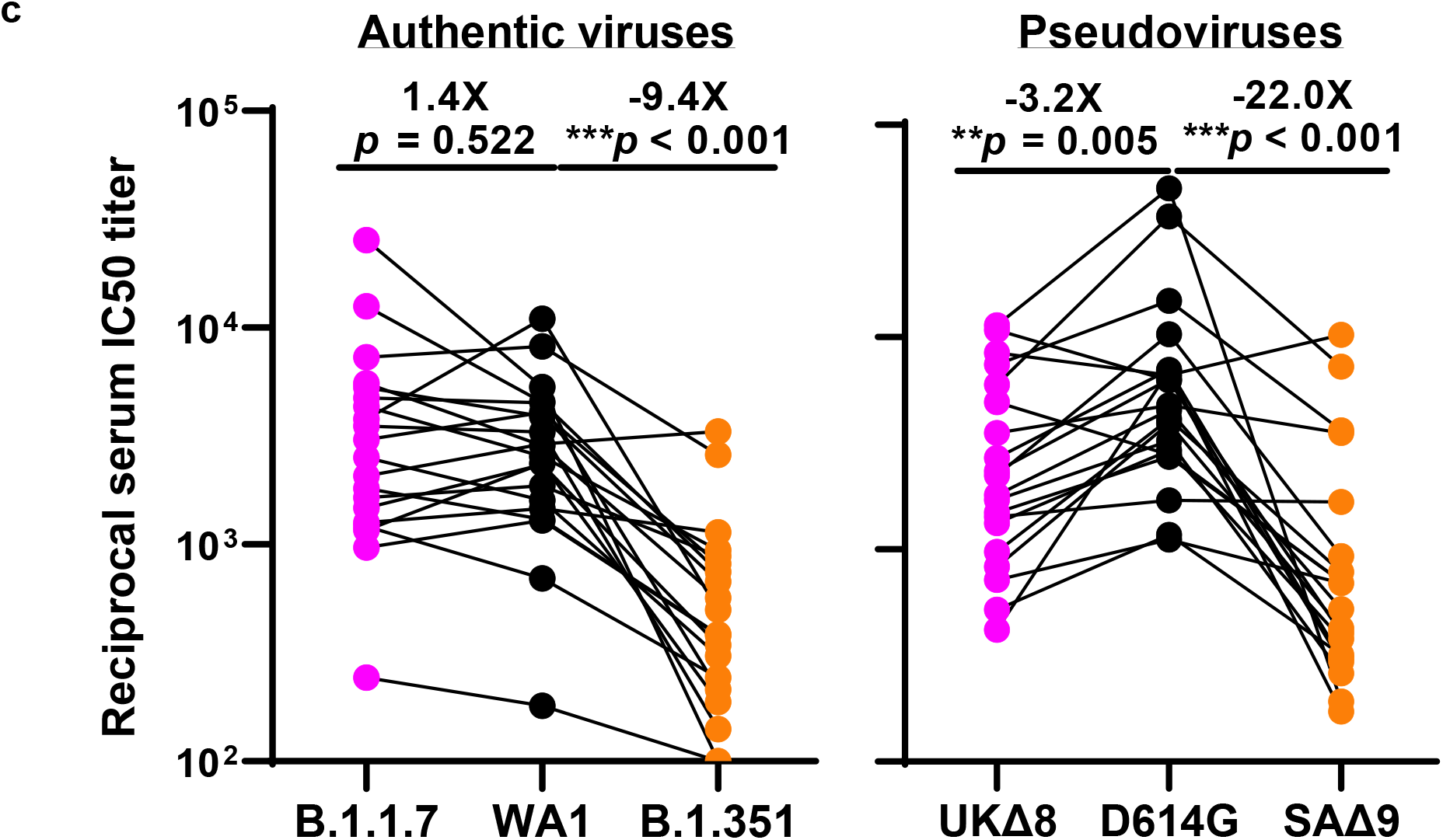
B.1.351 is more resistant to neutralization by convalescent plasma from patients. **a,** Neutralization results for 20 convalescent plasma samples (P1-P20) against B.1.1.7, B.1.351, and WT viruses. Data represent mean ± SEM of technical triplicates. **b,** Fold increase or decrease in neutralization IC50 of B.1.1.7 and B.1.351, as well as UKΔ8, SAΔ9, and single-mutation pseudoviruses, relative to the WT presented as a heatmap with darker colors implying greater change. **c,** Change in reciprocal plasma neutralization IC50 values of convalescent plasma against B.1.1.7 and B.1.351, as well as UKΔ8 and SAΔ9, relative to the WT. Mean fold changes in IC50 values relative to the WT are written above the *p* values. Statistical analysis was performed using a Wilcoxon matched-pairs signed rank test. Two-tailed p-values are reported.

Every plasma sample was also tested against each mutant pseudovirus, and those findings are shown in Extended Data Fig. 6 and summarized in Figs. 3b & 3c. Eight samples show >2.5-fold decrease in neutralizing activity against UKΔ8, in contrast to the results for B.1.1.7 neutralization. These discrepant results highlight our previous observation^20^ that pseudovirus neutralization does not always faithfully recapitulate live virus neutralization. The loss of plasma neutralizing activity against B.1.351 could be largely attributed to E484K (Fig. 3b), which has been shown to attenuate the neutralizing activity of convalescent sera^29^. Our findings here suggests that this RBM mutation is situated in an immunodominant epitope for most infected persons. It is also interesting to note that cases such as P7, P10, and P18 have neutralizing antibodies that are essentially unperturbed by the multitude of spike mutations found in these two new variants (Fig. 3b). A detailed analysis of their antibody repertoire against the viral spike could be informative.

## Vaccinee Sera

Sera were obtained from 12 participants of a Phase 1 clinical trial of Moderna SARS-Co-2 mRNA-1273 Vaccine^9^ conducted at the NIH. These volunteers received two immunizations with the vaccine (100 μg) on days 0 and 28, and blood was collected on day 43. Additional vaccinee sera were obtained from 10 individuals who received the Pfizer BNT162b2 Covid-19 Vaccine^10^ under emergency use authorization at the clinical dose on days 0 and 21. Blood was collected on day 28 or later.

Each vaccinee serum sample was assayed for neutralization against B.1.1.7, B.1.351, and WT viruses. Fig. 4a shows no loss of neutralizing activity against B.1.1.7, whereas every sample lost activity against B.1.351. These results are quantified and tabulated as fold increase or decrease in neutralization IC50 titers in Fig. 4b, and the extent of the decline in neutralization activity is more evident in Fig. 4c. Overall, the neutralizing activity against B.1.1.7 was essentially unchanged, but significantly lower against B.1.351 (12.4 fold, Moderna; 10.3 fold, Pfizer).

**Fig. 4.**
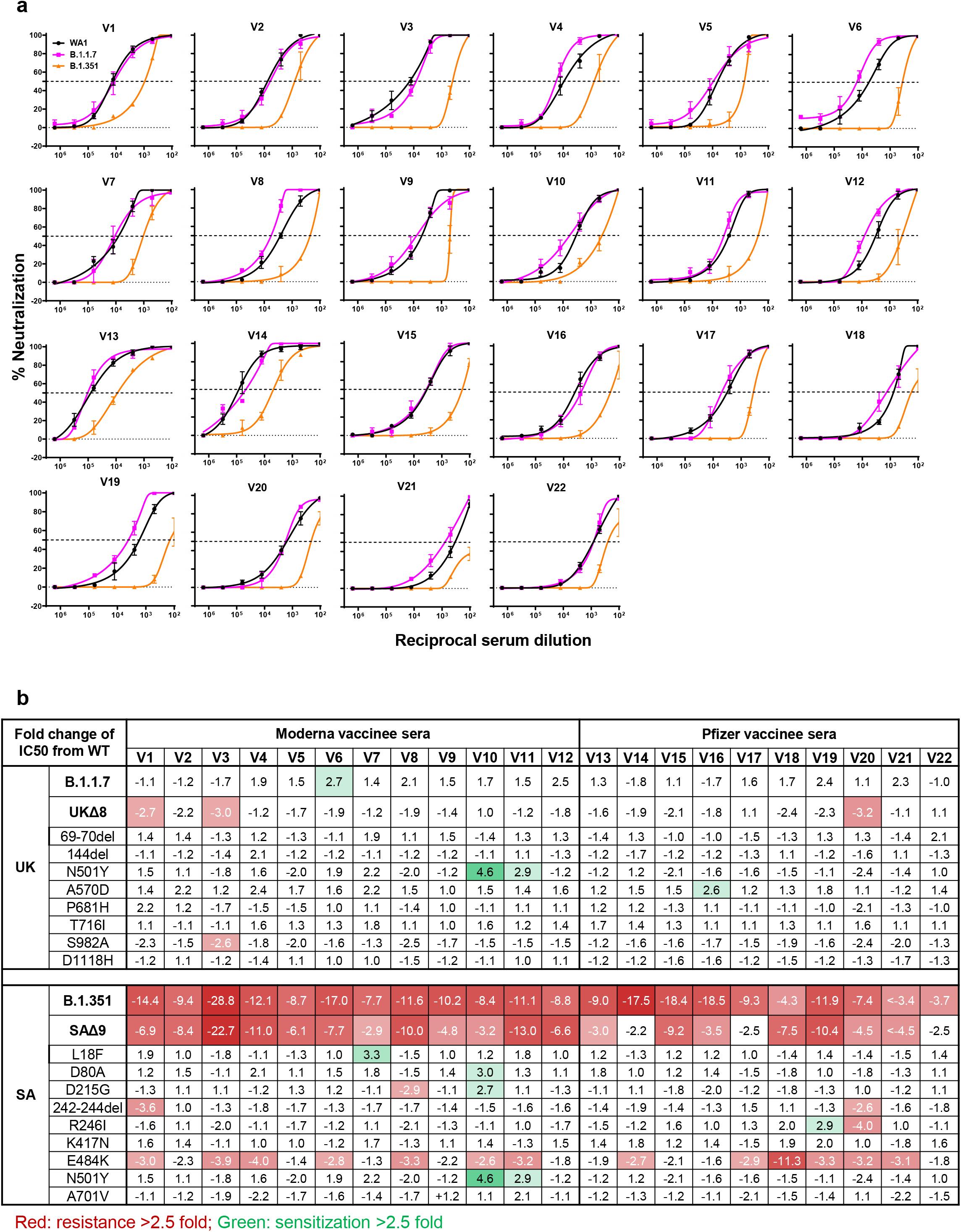
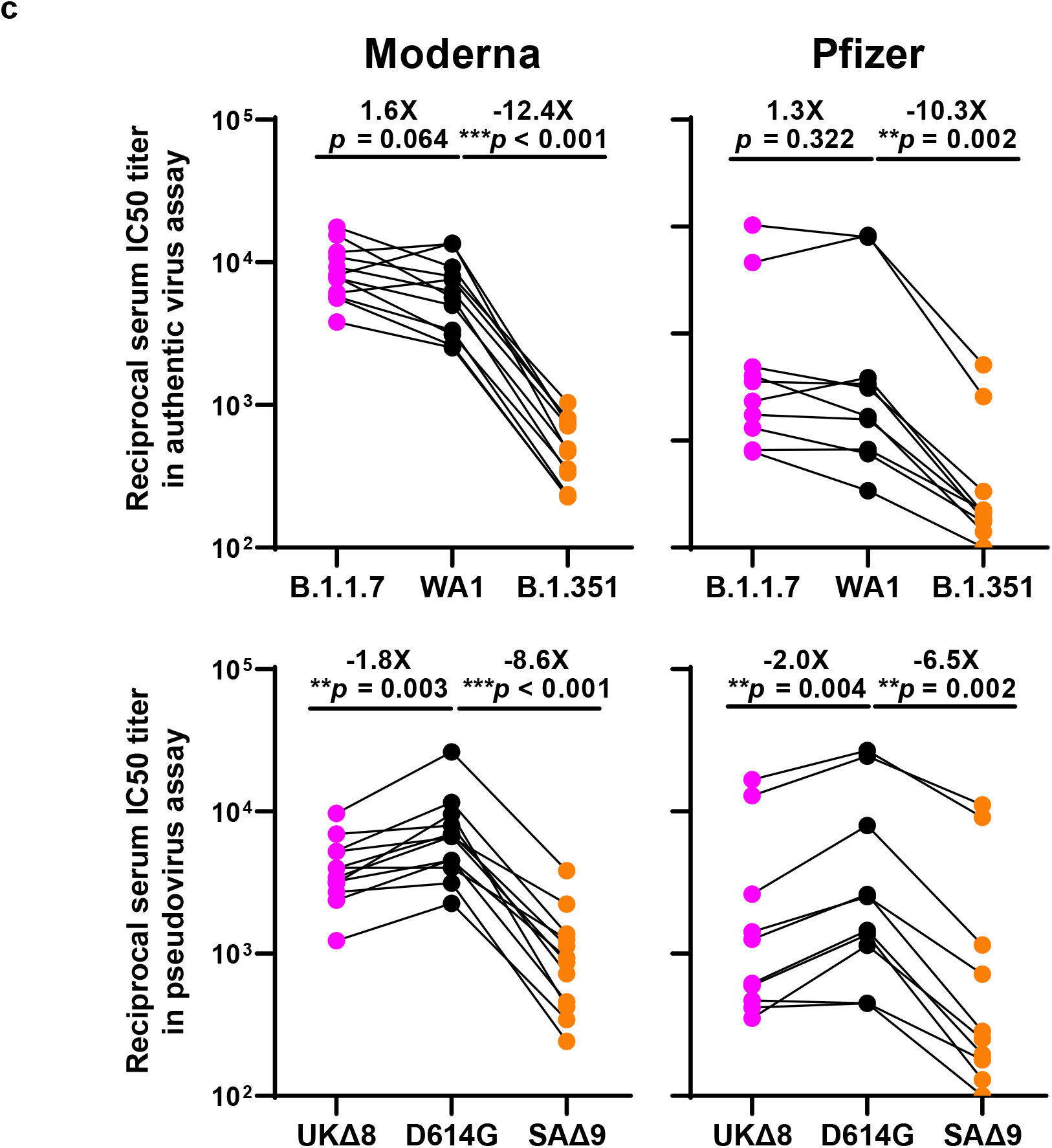
B.1.351 is more resistant to neutralization by vaccinee sera. **a,** Neutralization profiles for 22 serum samples obtained from persons who received SARS-CoV-2 vaccine made by Moderna (V1-V12) or Pfizer (V13-V22) against B.1.1.7, B.1.351, and WT viruses. Data are mean ± SEM of technical triplicates, and represent one of two independent experiments. **b,** Fold change in serum neutralization IC50 of B.1.1.7 and B.1.351, as well as UKΔ8, SAΔ9, and single-mutation pseudoviruses, relative to the WT, presented as a heatmap with darker colors implying greater change. **c,** Change in reciprocal serum IC50 values for Moderna and Pfizer vaccinees against B.1.1.7 and B.1.351, as well as UKΔ8 and SAΔ9, relative to the WT. Mean fold change in IC50 relative to the WT is written above the *p* values. Statistical analysis was performed using a Wilcoxon matched-pairs signed rank test. Two-tailed p-values are reported.

Every vaccinee serum was also tested against each mutant pseudovirus, and the results are presented in Extended Data Fig. 7 and summarized in Figs. 4b & 4c. No single mutation in B.1.1.7 has an appreciable impact on the neutralizing activity of vaccinee sera. The loss of neutralizing activity against SAΔ9 is largely consistent with the loss in B.1.351 live virus neutralization. A major contributor to the neutralization resistance of this variant virus appears to be E484K (Fig. 4b), indicating that this RBM mutation is situated in an immunodominant epitope recognized by all vaccinees studied.

## Discussion

Both SARS-CoV-2 variants B.1.1.7 and B.1.351 are raising concerns not only because of their increase transmissibility but also because of their extensive mutations in spike that could lead to antigenic changes detrimental to mAb therapies and vaccine protection. It is of equal concern that another variant known as P.1 or 501Y.V3 is increasing rapidly in Brazil and spreading far beyond^13,14^. P.1 contains three mutations (K417T, E484K, and N501Y) at the same RBD residues as B.1.351. Much of our findings on B.1.351 would likely be similar for this emergent variant. N501Y is shared among viruses in these three lineages; while this mutation may confer enhanced binding to ACE2^30^, its antigenic impact is limited to a few mAbs (Fig. 2c & Extended Data Fig. 4b), with no pronounced effects on the neutralizing activity of convalescent plasma or vaccinee sera (Figs. 3b & 4b), as others are reporting^31–33^.

Our findings have relevance to the use of mAb to treat or prevent SARS-CoV-2. Both B.1.1.7 and B.1.351 are resistant to neutralization by mAbs directed to the NTD supersite (Figs. 2c, 2d, & Extended Data Fig. 3b). More importantly, B.1.351 is resistant to a major group of potent mAbs that target the RBM, including three regimens authorized for emergency use (Fig. 2c). LY-CoV555 alone and in combination with CB6 are inactive against B.1.351, and the activity of REGN10933 is impaired (Fig. 2b) while its combination with REGN10987 retains much of the activity (Fig. 2e). Several other mAbs in development are similarly impaired (Figs. 2c, 2e, & Extended Data Fig. 4b) against this variant. Decisions on the use of these mAbs will depend heavily on the local prevalence of B.1.351 or variants with an E484K mutation, thus highlighting the importance of viral genomic surveillance worldwide and proactive development of next-generation antibody therapeutics, including combinations that target antigenically distinct epitopes.

Convalescent plasma from patients infected with SARS-CoV-2 from early in the pandemic show no significant change in neutralizing activity against B.1.1.7, but the diminution against B.1.351 is remarkable (Figs. 3b & 3c). This relative resistance is largely due to E484K, a mutation shared by B.1.351 and P.1^12–14^. Again, in areas where such viruses are common, one would have a concern about re-infection, as other studies are also suggesting^34,35^. This apprehension is heightened by the recent observation from the Novavax vaccine trial in South Africa that placebo recipients with prior SARS-CoV-2 infection were not protected against a subsequent exposure to B.1.351^36,37^. Even more disturbing is the situation in Manaus, Brazil where a second wave of infection due to P.1 is sweeping through a population that was already 76% seropositive due to prior infection in the Spring of 2020^38^.

As for the ramifications of our findings for the protective efficacy of current SARS-CoV-2 vaccines, the neutralizing activity of vaccinee sera against B.1.1.7 is largely intact and no adverse impact on current vaccines is expected (Fig. 4c), consistent with conclusions being reached by others^33,39,40^. On the other hand, the loss of 10.3-12.4 fold in activity against B.1.351 is larger than results being reported using mutant pseudoviruses^33,41,42^ or live virus^43^. Taken together, the overall findings are worrisome, particularly in light of recent reports that both Novavax and Johnson & Johnson vaccines showed a substantial drop in efficacy in South Africa^36,37,44^.

The recent emergence of B.1.1.7, B.1.351, and P.1 marks the beginning of SARS-CoV-2 antigenic drift. This conclusion is supported by data presented herein, illustrating how so many of these spike changes conferred resistance to antibody neutralization, and by studies reporting similar spike mutations selected by antibody pressure in vitro or in vivo^45–49^. Mutationally, this virus is traveling in a direction that could ultimately lead to escape from our current therapeutic and prophylactic interventions directed to the viral spike. If the rampant spread of the virus continues and more critical mutations accumulate, then we may be condemned to chasing after the evolving SARS-CoV-2 continually, as we have long done for influenza virus. Such considerations require that we stop virus transmission as quickly as is feasible, by redoubling our mitigation measures and by expediting vaccine rollout.

