## Supplementary material for "Antibody Resistance of SARS-CoV-2 Variants B.1.351 and B.1.1.7": Methods and Extended Data

**Patients and vaccinees.** Plasma samples were obtained from patients (age 34-79; mean 54) convalescing from documented SARS-CoV-2 infection approximately one month after recovery or later. These cases were enrolled into an observational cohort study of convalescent patients followed at the Columbia University Irving Medical Center starting in the Spring of 2020. The study protocol was approved by the Institutional Review Board (IRB), and all participants provided written informed consent. From their documented clinical profiles, plasma samples from ten with severe Covid-19 were selected, along with plasma from 10 with non-severe infection, for this study. Sera were obtained from 12 participants in a Phase 1 clinical trial of Moderna SARS-CoV-2 mRNA-1273 Vaccine conducted at the NIH, under an IRB-approved protocol. Sera were also obtained from 10 individuals followed in an IRB-approved protocol to assess immunological responses to SARS-CoV-2 who received the Pfizer BNT162b2 Covid-19 Vaccine as a part of the emergency use authorization

**Monoclonal antibodies.** Monoclonal antibodies tested in this study were constructed and produced at Columbia University as previously described[20], except REGN10933, REGN10987, REGN10985, COV2-2196, and COV2-2130 were provided by Regeneron Pharmaceuticals, Inc., Brii-196 and Brii-198 were provided by Brii Biosciences, and CB6 was provided by B.Z. and P.D.K.

**Authentic SARS-CoV-2 Microplate Neutralization.** The SARS-CoV-2 viruses USA-WA1/2020 (WA1), USA/CA_CDC_5574/2020 (B.1.1.7), and hCoV-19/South Africa/KRISP-EC-K005321/2020 (B.1.351) were obtained from BEI Resources (NIAID, NIH) and propagated for one passage using Vero-E6 cells. Virus infectious titer was determined by an end-point dilution and cytopathic effect (CPE) assay on Vero-E6 cells as described previously[20].

An end-point dilution microplate neutralization assay was performed to measure the neutralization activity of convalescent plasma samples, vaccinee sera, and purified mAbs. In brief, plasma and serum samples were subjected to successive 5-fold dilutions starting from 1:100. Similarly, most mAbs were serially diluted (5-fold dilutions) starting at 10 μg/mL. Some clinical antibodies were tested from starting concentrations of 1 μg/mL. Triplicates of each dilution were incubated with SARS-CoV-2 at an MOI of 0.1 in EMEM with 7.5% inactivated fetal calf serum (FCS) for 1 hour at 37$^{o}$C. Post incubation, the virus-antibody mixture was transferred onto a monolayer of Vero-E6 cells grown overnight. The cells were incubated with the mixture for ~70 hours. Cytopathic effect (CPE) of viral infection was visually scored for each well in a blinded fashion by two independent observers. The results were then converted into percentage neutralization at a given sample dilution or mAb concentration, and the averages ± SEM were plotted using a five-parameter dose-response curve in GraphPad Prism v8.4.

**Construction and production of variant pseudoviruses.** The original pCMV3-SARS-CoV-2-spike plasmid was kindly provided by Dr. Peihui Wang of Shandong University in China. Plasmids encoding for D614G variant, all the single-mutation variants found in B.1.1.7 or B.1.351, 8-mutation-combination variant (UKΔ8) and 9-mutation-combination variant (SAΔ9) were generated by Quikchange II XL site-directed mutagenesis kit (Agilent). Recombinant Indiana VSV (rVSV) expressing different SARS-CoV-2 spike variants were generated as previously described[20,21]. HEK293T cells were grown to 80% confluency before transfection with the spike gene using Lipofectamine 3000 (Invitrogen). Cells were cultured overnight at 37°C with 5% $CO_2$, and VSV-G pseudo-typed ΔG-luciferase (G*ΔG-luciferase, Kerafast) was used to infect the cells in DMEM at an MOI of 3 for 2 hours before washing the cells with 1X DPBS three times. The next day, the transfection supernatant was harvested and clarified by centrifugation at 300 g for 10 min. Each viral stock was then incubated with 20% I1 hybridoma (anti-VSV-G, ATCC: CRL-2700) supernatant for 1 hour at 37°C to neutralize contaminating VSV-G pseudo-typed ΔG-luciferase virus before measuring titers and making aliquots to be stored at −80°C.

**Pseudovirus neutralization assays.** Neutralization assays were performed by incubating pseudoviruses with serial dilutions of mAbs or heat-inactivated plasma or sera, and scored by the reduction in luciferase gene expression[20,21]. In brief, Vero E6 cells were seeded in a 96-well plate at a concentration of $2 \times 10^4$ cells per well. Pseudoviruses were incubated the next day with serial dilutions of the test samples in triplicate for 30 minutes at 37°C. The mixture was added to cultured cells and incubated for an additional 24 hours. The luminescence was measured by Luciferase Assay System (Promega). $IC_{50}$ was defined as the dilution at which the relative light units were reduced by 50% compared to the virus control wells (virus + cells) after subtraction of the background in the control groups with cells only. The $IC_{50}$ values were calculated using non-linear regression in GraphPad Prism.

**Reporting summary.** Further information on research design is available in the Nature Research Reporting Summary linked to this paper.

**Data availability.** Materials used in this study will be made available but may require execution of a materials transfer agreement. Source data are provided herein.

**Acknowledgements.** We thank Stephen Goff and Brandon DeKosky for helpful discussions. This study was supported by funding from Andrew & Peggy Cherng, Samuel Yin, Barbara Picower and the JPB Foundation, Brii Biosciences, Roger & David Wu, and the Bill and Melinda Gates Foundation.

**Author contributions.** The study was conceptualized by D.D.H. The experiments were principally carried out by P.W., M.N., Y.H., L.L., and S.I. with able assistance from M.W., J.Y. Structural interpretations were made by Y.G., Z.S., L.S., and P.D.K. B.Z., P.D.K., and C.A.K. provided mAbs. J.Y.C. and M.T.Y. provided plasma from convalescent patients. B.S.G. and J.R.M. provided sera from participants in the Moderna vaccine trial; J.Y.C. and M.S. provided sera from health care workers immunized with the Pfizer vaccine. Y.H. and Y.L. helped to supervise the study. The manuscript was written by D.D.H. with editing by P.W., P.D.K., L.S., Y.L., and reviewed, commented, and approved by all the authors.

**Competing interests:** P.W., L.L, J.Y., M.N., Y.H., and D.D.H. are inventors on a provisional patent application on mAbs to SARS-CoV-2. D.D.H. is a member of the scientific advisory board of Brii Biosciences, which also has provided a grant to Columbia University to support this and other studies on SARS-CoV-2.

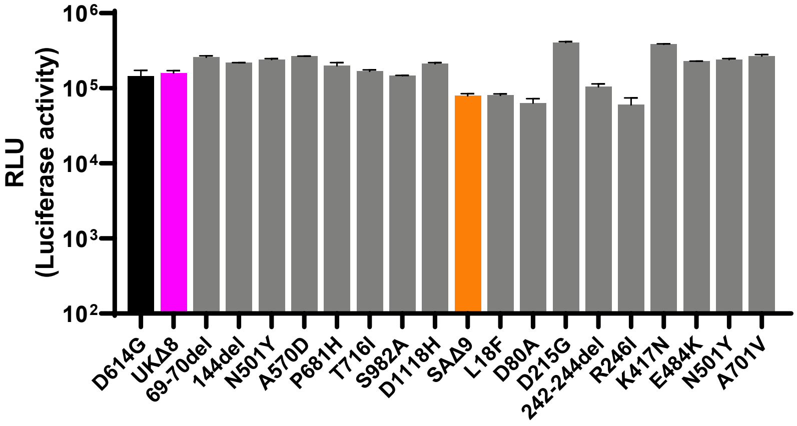


**Extended Data Fig. 1 | Titers of WT (D614G) and the 18 mutant SARS-CoV-2 pseudoviruses.** VSV-based pseudoviruses were generated[18,19] and viral particles were quantified and normalized by VSV nucleocapsid protein by western blot. Equal amount of each pseudovirus was then used to infect Vero E6 cells and relative luciferase unit (RLU) was measured 16 hrs later.

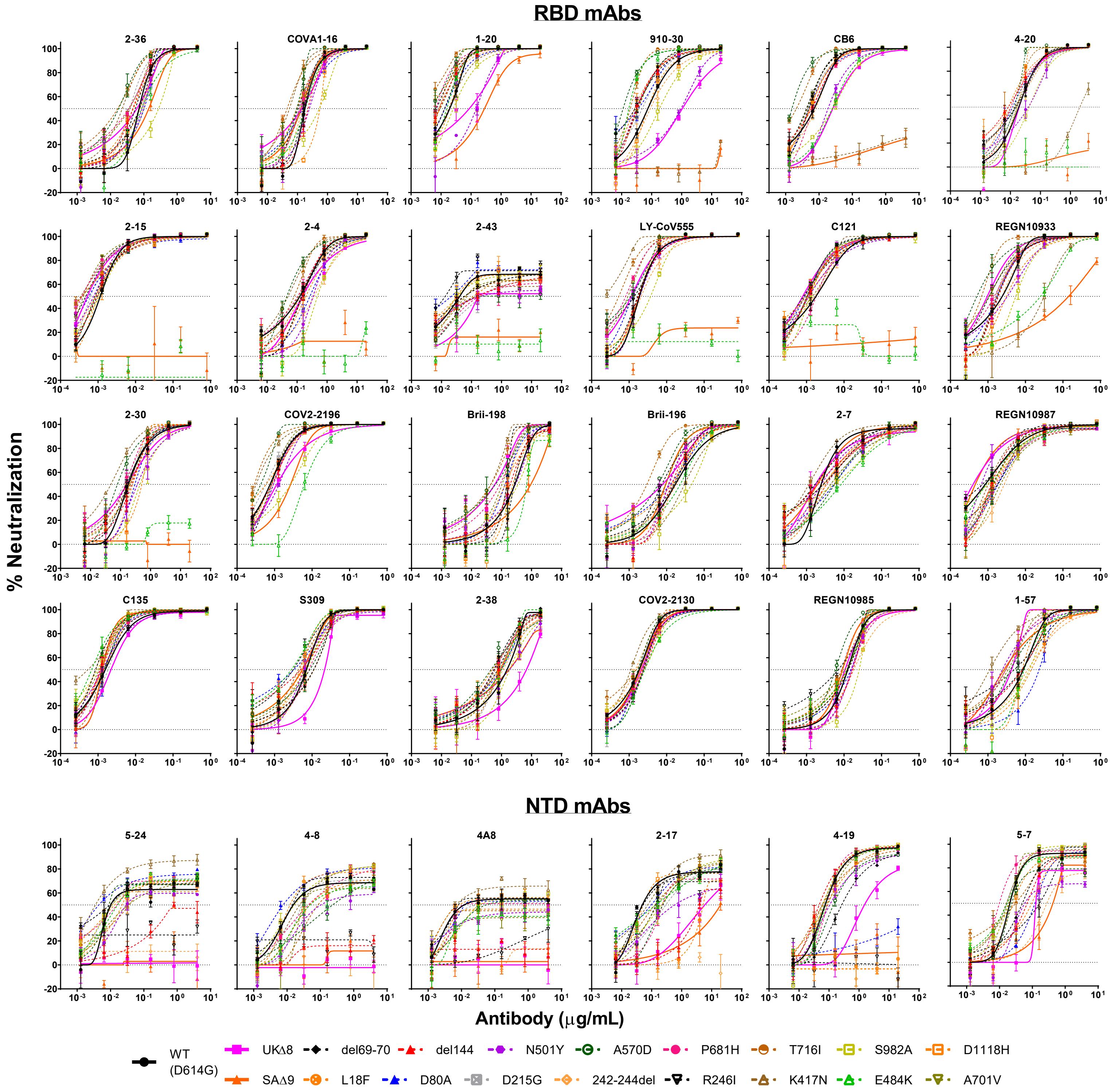


**Extended Data Fig. 2 | Neutralization profiles of mAbs against WT, UKΔ8, and SAΔ9, as well as single-mutation pseudoviruses.** Data represent mean ± SEM of technical triplicates.

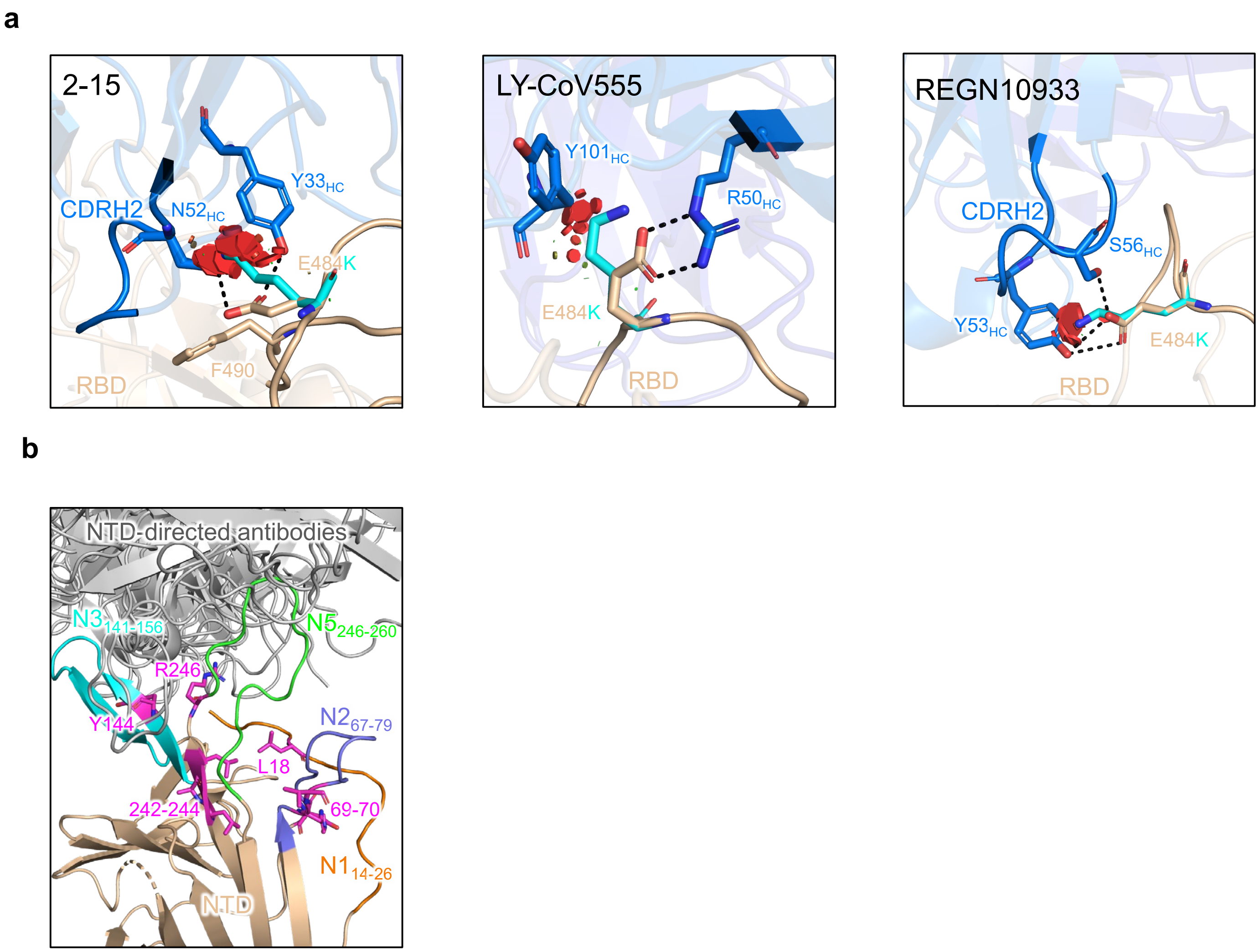


**Extended Data Fig. 3 | Structural explanations on how critical mutations affect mAb activity. a,** E484 forms hydrogen bonds with mAbs that target RBM. Mutation E484K causes not only steric clashes but also a charge change at antibody binding sites, and thus abolishes binding by these RBM-directed mAbs. Steric clashes are shown by red plates. **b,** Mutations at or near the NTD antigenic supersite – comprised of loops N1, N3, and N5 – that is recognized by many potent NTD-directed mAbs.

**a**

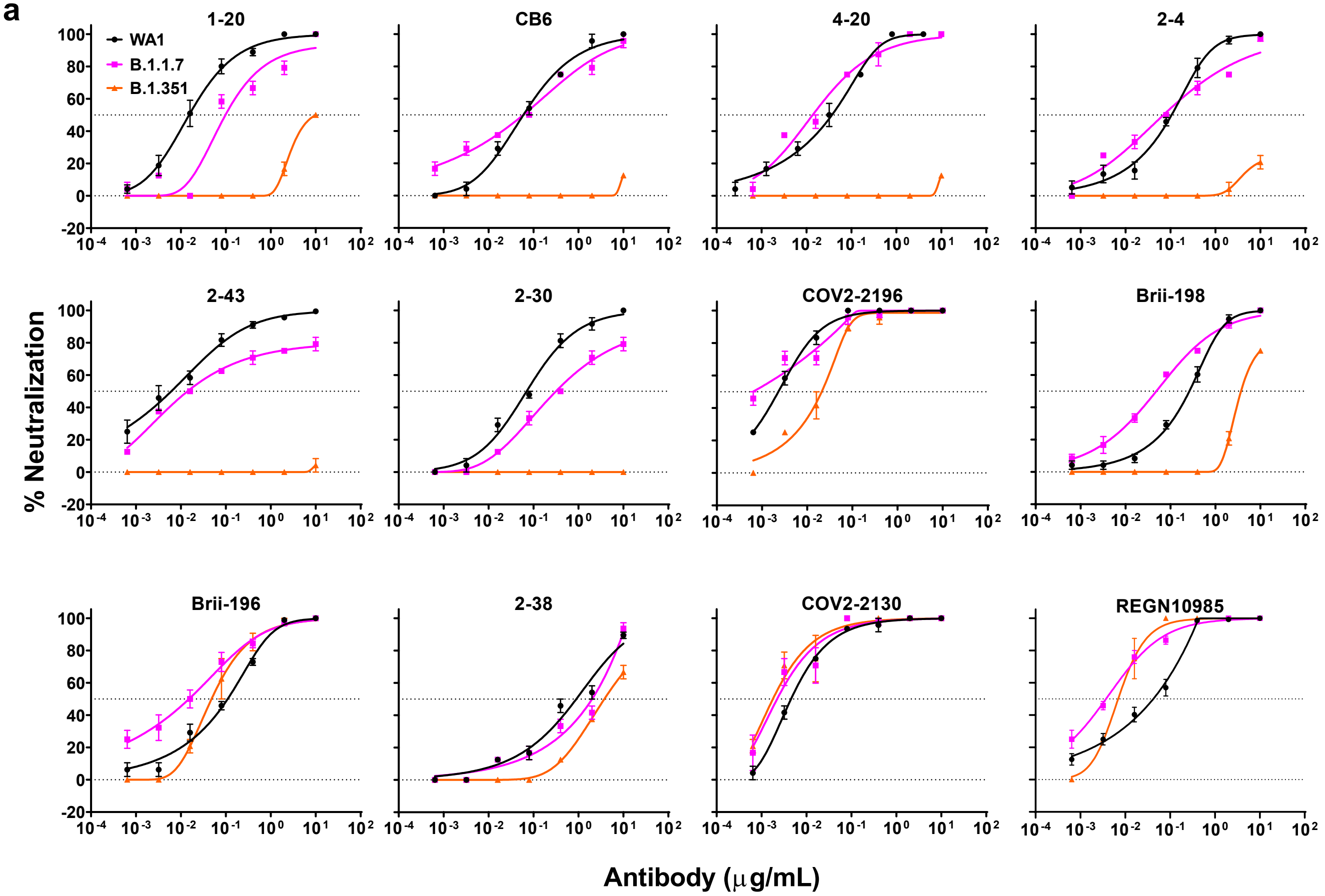


**b**

| Fold change of IC50 from WT | | RBD-directed mAbs | | | | | | | | | | | |
|---|---|---|---|---|---|---|---|---|---|---|---|---|---|
| | | 1-20 | CB6 | 4-20 | 2-4 | 2-43 | 2-30 | COV2-2196 | Brii-198 | Brii-196 | 2-38 | COV2-2130 | REGN10985 |
| UK | **B.1.1.7** | -5.6 | 1.4 | 2.4 | 1.3 | -2.3 | -4.4 | 3.0 | 5.6 | 7.3 | -1.8 | 3.0 | 10.5 |
| | **UKΔ8** | -4.5 | -3.6 | -1.0 | -1.2 | -4.1 | 1.1 | -1.5 | 4.6 | 2.2 | -4.4 | -1.1 | 1.2 |
| | 69-70del | 1.5 | 1.5 | 1.2 | -1.2 | -1.4 | 1.1 | 1.0 | 1.1 | 1.1 | 2.8 | -1.3 | -1.0 |
| | 144del | 2.2 | 1.2 | 2.2 | -1.2 | -3.7 | -1.3 | -1.1 | 1.4 | 1.2 | 1.6 | -1.0 | -1.4 |
| | N501Y | -7.7 | -2.6 | -2.1 | -2.8 | -4.8 | -2.0 | -1.7 | -1.0 | 1.5 | -1.2 | -1.3 | -1.3 |
| | A570D | 4.6 | 4.6 | 1.4 | 2.7 | -5.3 | 2.3 | 1.8 | 4.4 | 2.9 | 3.4 | 1.1 | 2.1 |
| | P681H | 2.9 | 1.1 | 1.8 | 1.2 | -4.5 | 1.3 | -1.2 | 3.1 | 1.5 | 2.7 | -1.4 | -1.3 |
| | T716I | 10.0 | 3.4 | 1.9 | 1.5 | -1.5 | -1.1 | 2.6 | 6.3 | 5.7 | 1.7 | 1.2 | 1.4 |
| | S982A | -1.3 | -3.6 | -2.2 | -3.9 | 1.2 | -2.4 | -3.6 | -1.3 | -2.7 | -1.2 | -1.4 | -2.0 |
| | D1118H | 1.0 | 1.4 | 1.1 | -1.6 | 1.2 | -2.8 | 1.3 | 1.2 | 1.2 | 1.8 | -1.2 | 1.0 |
| SA | **B.1.351** | -667.0 | <-1000 | <-1000 | <-1000 | <-1000 | <-1000 | -6.3 | -14.6 | 2.2 | -3.5 | 3.0 | 5.3 |
| | **SAΔ9** | -14.4 | <-1000 | <-1000 | <-1000 | <-1000 | <-1000 | -3.5 | -3.4 | 1.8 | -1.2 | -1.0 | 1.4 |
| | L18F | 2.2 | 1.3 | 2.2 | -1.0 | -2.8 | -1.1 | 1.0 | 2.1 | 1.1 | 3.1 | -1.2 | 1.1 |
| | D80A | 1.4 | 1.4 | 1.6 | -2.4 | 1.9 | -1.1 | -1.2 | 2.1 | 1.4 | 1.6 | -1.0 | -1.2 |
| | D215G | 2.4 | 1.6 | 1.3 | -1.4 | -5.6 | 1.4 | 1.3 | 1.7 | 1.0 | 1.5 | 1.1 | 1.4 |
| | 242-244del | 5.1 | -1.1 | 1.5 | -2.1 | 1.2 | 1.1 | 1.0 | 1.7 | -1.1 | 1.4 | 1.3 | -1.4 |
| | R246I | 2.8 | 1.7 | 1.3 | -1.1 | 3.0 | 1.2 | -1.1 | 1.9 | 2.3 | 2.0 | -1.2 | 1.8 |
| | K417N | 1.1 | <-1000 | -108.3 | 2.5 | -1.7 | 3.8 | 2.4 | 6.3 | -1.8 | 2.5 | 1.7 | 1.1 |
| | E484K | 1.3 | -3.5 | <-1000 | <-1000 | <-1000 | <-1000 | -8.1 | -2.5 | -1.4 | 1.5 | -1.4 | 1.0 |
| | N501Y | -7.7 | -2.6 | -2.1 | -2.8 | -4.8 | -2.0 | -1.7 | -1.0 | 1.5 | -1.2 | -1.3 | -1.3 |
| | A701V | 2.1 | 1.6 | 1.1 | 1.1 | -1.6 | -1.2 | -1.1 | 2.7 | 1.8 | 1.7 | -1.0 | 1.2 |

Red: resistance >3 fold; Green: sensitization >3 fold

**Extended Data Fig. 4 | Neutralization susceptibility of UK and SA variants to additional SARS-CoV-2 RBD-directed mAbs. a,** Neutralization of B.1.1.7, B.1.351, and WT viruses by additional RBD-directed mAbs. Data represent mean ± SEM of technical triplicates. **b,** Fold increase or decrease in IC50 of neutralizing mAbs against B.1.1.7 and B.1.351, as well as mutant pseudoviruses, relative to WT.

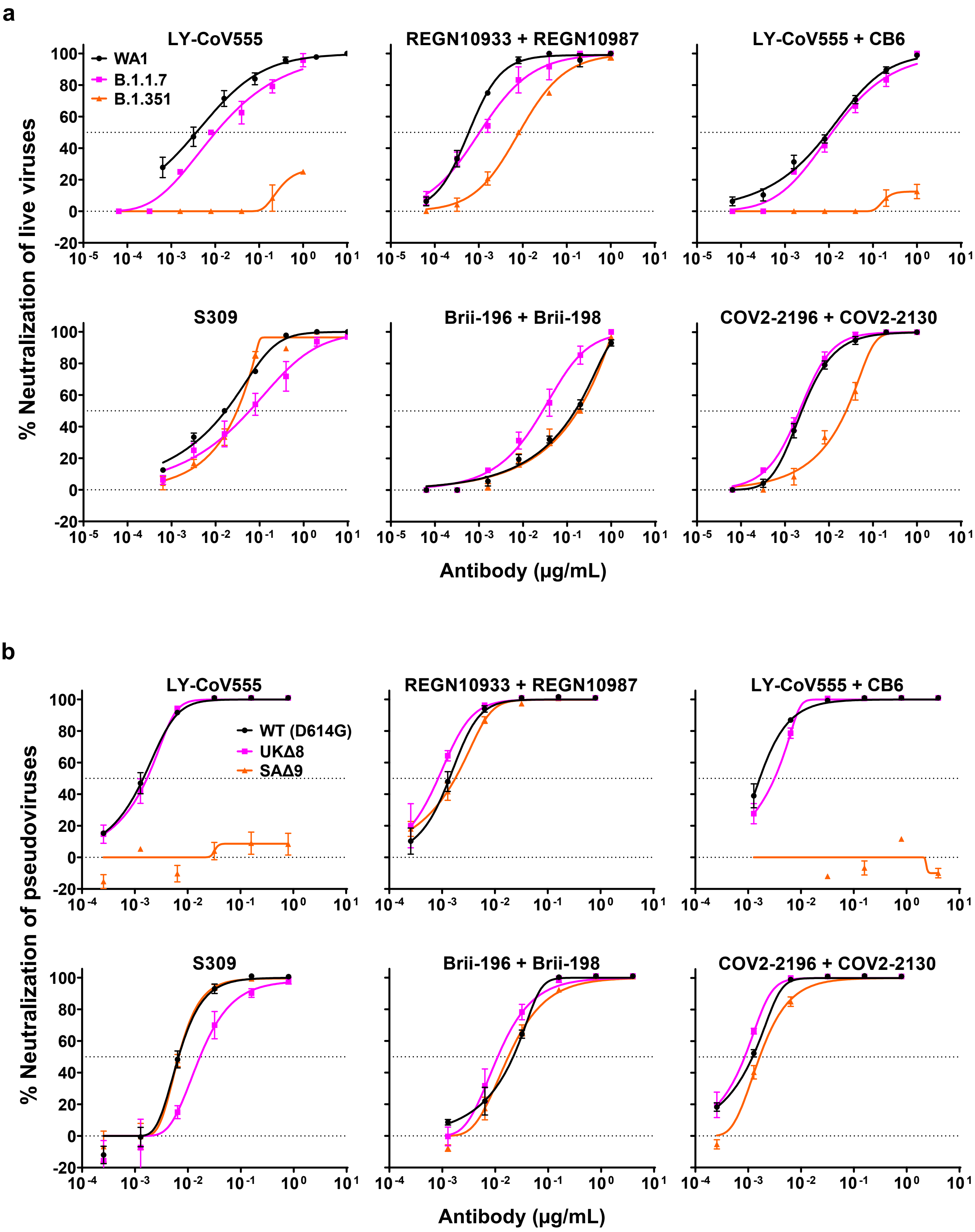


**Extended Data Fig. 5 | Neutralization profiles of authorized or investigational therapeutic mAbs against WT (WA1), B.1.1.7, and B.1.351 live viruses (a), and against WT, UKΔ8, and SAΔ9 pseudoviruses (b).** Data represent mean ± SEM of technical triplicates.

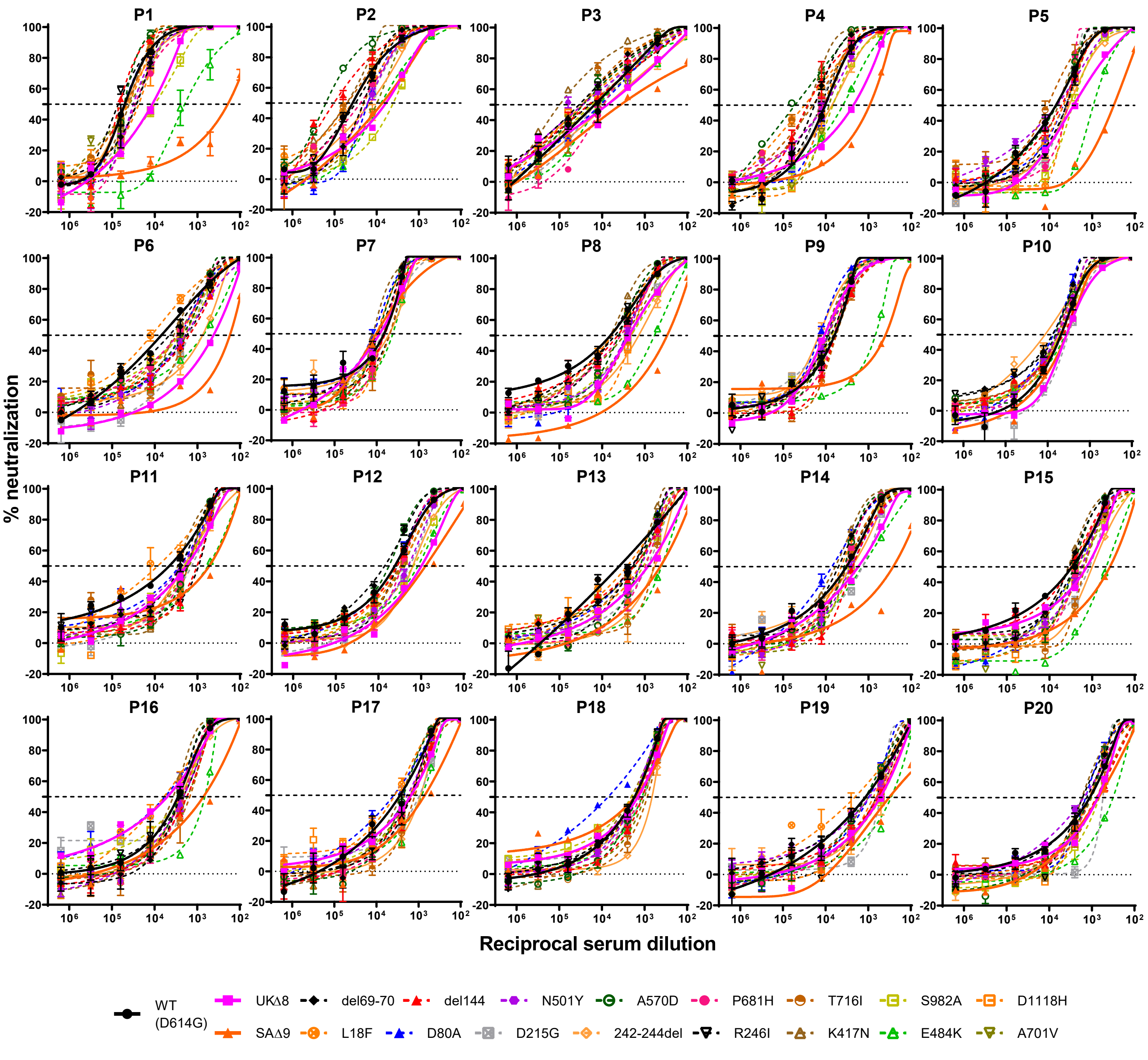


**Extended Data Fig. 6 | Neutralization profiles of 20 convalescent patient plasma against WT, UKΔ8, SAΔ9, and single-mutation pseudoviruses.** Data represent mean ± SEM of technical triplicates.

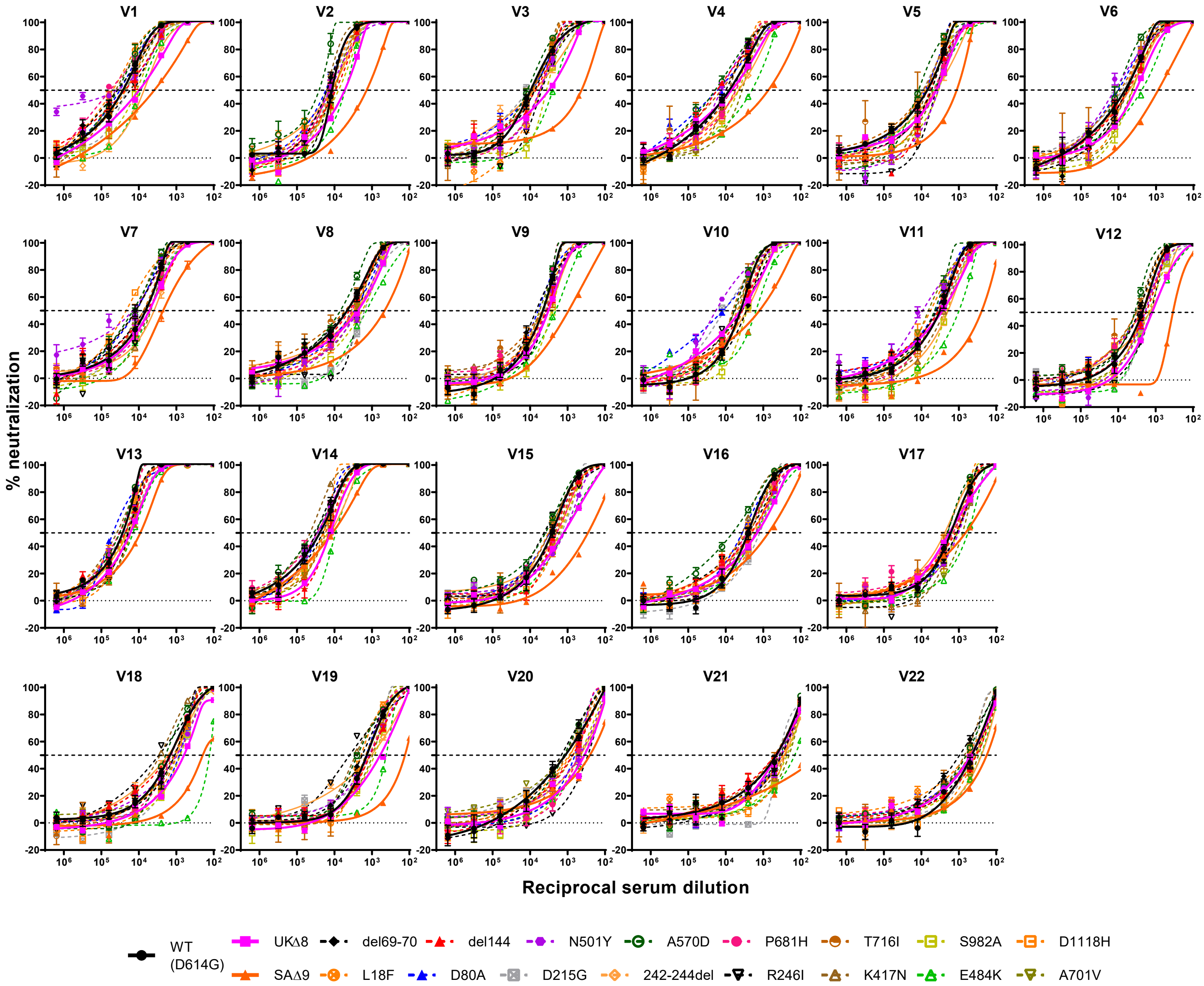


**Extended Data Fig. 7 | Neutralization profiles of vaccinee sera against WT, UKΔ8, SAΔ9, and single-mutation pseudoviruses.** 12 sera from Moderna vaccinees (V1-V12) and 10 sera from Pfizer vaccinees (V13-V22) were tested. Data represent mean ± SEM of technical triplicates.
